# Divergent sex differences in functional brain connectivity networks in excessively drinking C57BL/6J mice

**DOI:** 10.1101/2021.05.19.444869

**Authors:** Solal Bloch, Jennifer A. Rinker, Alex C.W. Smith, Priyattam J. Shiromani, Damian G. Wheeler, Ricardo Azevedo, Sunil Gandhi, Michaela Hoffman, Patrick J. Mulholland

## Abstract

Individuals with alcohol use disorder continue to drink in excess despite the health and societal consequences, and the rate of problematic drinking and alcohol-related harms is increased in women. Clinical imaging studies report widespread adaptations in brain structure after chronic, heavy drinking, and alcohol-related cues enhance brain reactivity in reward-related regions. In rodents, alcohol drinking induces expression of the immediate early gene c-Fos, which can be a marker of cellular activity, across multiple brain regions. Recent evidence also suggests that abstinence from chronic intermittent alcohol exposure can produce mesoscale changes in c-Fos expression. However, there is a substantial gap in our understanding of how excessive drinking affects functional connectivity networks to influence alcohol-seeking behaviors. For this study, male and female C57BL/6J mice were given access to either water or a choice between water and ethanol in the intermittent access drinking model for 4 weeks. After a short-access drinking session, whole brains from high alcohol drinking male and female mice and water drinking controls were then subjected to c-Fos immunolabeling, iDISCO+ clearing, light sheet imaging, and whole-brain c-Fos mapping. Correlation matrices were then generated and graph theoretical statistical approaches were used to determine changes in functional connectivity across sex and drinking condition. We observed robust sex differences in the network of c-Fos+ cells in water drinking mice, and excessive alcohol drinking produce divergent and robust changes in functional network connectivity in male and female mice. In addition, these analyses identified novel hub regions in excessively drinking mice that were unique for each sex. In conclusion, the whole-brain c-Fos mapping analysis identified sex difference in functional network connectivity and unique and understudied regions that may play a critical role in controlling excessive ethanol drinking in male and female mice.

---

Alcohol use disorder (AUD) is a relapsing disease characterized by intense craving for alcohol and high relapse rates despite the known negative consequences (e.g., cardiovascular and liver disease, cancers) of continued drinking. In fact, >5% of all worldwide deaths (more than 3 million) each year are caused by excessive alcohol drinking (WHO, 2018). In the United States, per capita consumption of alcohol and binge drinking increased over the last two decades (Grucza et al., 2018; Martinez et al., 2019), as did alcohol-related deaths (White et al., 2020). Although the rate of AUD is higher in men (7.6% vs 4.1%), problematic alcohol use and alcohol-related harms are increasing sharply in women (Slade et al., 2016). The inability of individuals with AUD to control their drinking and reduce the long-term consequences of excessive consumption highlights the need to define the mechanisms and neurocircuits underlying compulsive alcohol use.

Structurally, there are gray matter changes throughout the brain, including core regions involved in reward processing, in individuals with AUD (Buhler and Mann, 2011; Shim et al., 2019). Additionally, there are widespread reductions in fractional anisotropy, a method to assess white matter microstructure, in AUD (Fortier et al., 2014; Fritz et al., 2019). While there is a decrease in neural activity as measured by fMRI and BOLD signals in AUD individuals under basal conditions (Buhler and Mann, 2011; Fritz et al., 2019; Kalivas and Volkow, 2005), there is enhanced reactivity to alcohol-related cues in regions thought to drive craving and relapse (Bracht et al., 2021; Buhler and Mann, 2011; Fritz et al., 2019). In the rodent literature, studies have shown that voluntary alcohol drinking induced expression of c-Fos, an immediate early gene, in a number of brain regions (Burnham and Thiele, 2017; Li et al., 2010; Ryabinin et al., 2003; Sajja and Rahman, 2013; Smith et al., 2020), most of which have been the target of extensive analysis. Importantly, a study showed that manipulating activated cells (i.e., c-Fos+) in the infralimbic cortex can have a strong influence over controlling excessive alcohol-seeking behaviors (Pfarr et al., 2015). More recently, evidence also suggests that abstinence from chronic intermittent alcohol exposure can change large-scale network activity as measured by whole-brain c-Fos expression (Kimbrough et al., 2020). However, there is a substantial gap in our understanding of how excessive drinking affects whole-brain functional connectivity networks to influence alcohol-seeking behaviors.

Recent technological advances in tissue clearing, light sheet imaging, and network analysis have rapidly gained interest in the neuroscience field. Brain clearing techniques, such as iDISCO+ (Renier et al., 2016), that allow for efficient deep tissue immunolabeling of whole mouse brains and refractive index matching improves the resolution of rapid volumetric imaging by minimizing light scattering and absorption. Computational analysis of large scale brain imaging and immediate early gene mapping have yielded excellent correspondence between functional measurements and anatomical connections (Honey et al., 2010; Wheeler et al., 2013). Despite these notable state-of-the-art methods and data analysis tools, whole-brain light sheet imaging has not been widely applied to study systems-level pathological mechanisms of alcohol and substance use disorders (Simpson et al., 2021). Thus, the purpose of this study was to characterize whole-brain c-Fos activity mapping in male and female mice that were drinking in an intermittent access to alcohol model that produces high amounts of intake (Cannady et al., 2020; Hwa et al., 2011; Rinker et al., 2017).

## Materials and Methods

### Animals

Male and female C57BL/6J mice were obtained from Jackson Laboratories (Bar Harbor, ME; https://www.jax.org/strain/00064) at 7 weeks of age. They were group-housed (4/cage) and allowed to acclimatize to the colony room for at least one week in a temperature and humidity-controlled AAALAC-approved facility. Animals were maintained on a reverse 12-h light/dark cycle with lights off at 09:00 am and had ad libitum access to food and water. All animals were treated in strict accordance with the NIH Guide for the Care and Use of Laboratory Animals and all experimental methods were approved by the Medical University of South Carolina’s Institutional Animal Care and Use Committee.

### Two-bottle choice intermittent ethanol access

After acclimatization, mice were housed individually and were given 24 h intermittent access to alcohol (IAA; 20% v/v) and water from 9 am to 9 am with 24 or 48 h between drinking sessions (Mondays, Wednesdays, and Fridays) (Bloch et al., 2020; Cannady et al., 2020; Rinker et al., 2017; Zamudio et al., 2020). Mice were subjected to the IAA model for 4 weeks with 3 drinking sessions/week. The location of ethanol and water bottles was alternated on each drinking session. All groups received two water bottles on intervening days. Procedures were identical in age-matched control mice except mice were given access to two water bottles during drinking sessions. Mice were sacrificed 30 min following a final 2 h drinking session and brains were extracted and prepared for whole-brain c-Fos mapping. Ethanol preference was calculated from the amount of ethanol consumed as a percentage of the total amount of fluid (ethanol + water) consumed during each drinking session.

### iDISCO+ Brain Clearing and c-Fos Immunolabeling

Following the drinking session, mice were perfused with 20 mL of PBS followed by 20 mL 4% PFA in PBS and post-fixed overnight at 4◻C. Brains were then dehydrated with methanol, bleached in 5% H_2_O_2_ in methanol before permeabilization, blocking, and incubation with primary and secondary antibodies following routine procedures for iDISCO+ labeling (Renier et al., 2016). Brains were then dehydrated and cleared with DCM and dibenzyl ether for refractive index matching. The polyclonal rabbit c-Fos primary antibody (Synaptic Systems, Catalog # 226003) and the donkey anti-rabbit Alexa Fluor 568 secondary antibody (ThermoFisher Scientific, Catalog # A10042) were used at a 1:300 dilution.

### Light Sheet Imaging and c-Fos Mapping

After immunolabeling and clearing, whole brains were imaged using a Zeiss Z.1 light sheet microscope (Zeiss Microscopy) and 5× objective (0.16 NA) fitted for the Mesoscale imaging chamber (Translucence Biosystems) with 1.28 × 1.28 × 5 µm resolution. The 488 nm laser was used to acquire the autofluorescence channel and the 561 laser was used for acquisition of c-Fos expression. Laser power and exposure were matched across samples. Images were stitched using Stitcher (Bitplane) software, and converted to TIF format using custom Python scripts. c-Fos detection and alignment of cell coordinates to the Allen Reference Atlas (25 µm resolution) was performed using BrainQuant3D software that was designed for acquisition using a Zeiss Z.1 light sheet microscope (Schneider et al., 2019). Cell counts for each region was normalized to region volume before further analysis.

### Network Analysis

Whole-brain c-Fos mapping data were analyzed using a multifaceted approach. First, standard functional connectivity matrices using Pearson correlations were calculated for each treatment group (Cruces-Solis et al., 2020; Kimbrough et al., 2020; Vetere et al., 2017). Correlation matrices were organized into 12 major divisions of gray matter, following the Allen Reference Atlas ontology according to the Common Coordinate Framework, version 3 (Wang et al., 2020): cerebellum, cortical subplate, hippocampal formation, hypothalamus, isocortex, medulla, midbrain, olfactory areas, pallidum, pons, striatum, and thalamus. For these analyses, subdivisions of individual regions and layer-specific subregions (e.g., prelimbic, layers I, II/III, V, VIa, VIb) were excluded, yielding 306 regions that were included in these analyses. Correlation coefficients were averaged within major divisions and analyzed across treatment groups and sex.

Theoretical and empirical data suggest that the mammalian brain is organized into a network with small-world architecture (Bullmore and Sporns, 2009; Rubinov and Sporns, 2010; Wheeler et al., 2013), where brain regions in close proximity to each other interact in an effort to reduce metabolic demands (Bullmore and Sporns, 2009; Farahani et al., 2019). However, there are a small number of long-range connections that are proposed to accelerate neural transmission across local networks. The application of Graph Theory to the neuroscience field allows researchers to construct and understand changes in local brain networks and long-range connections in health and disease (Sporns, 2018). To understand how excessive drinking affects properties of the network, we analyzed quantifiable measures of the clustering analysis (Rubinov and Sporns, 2010). These measures of the complex networks include assessments of functional integration (e.g., characteristic path length), segregation (e.g., transitivity), and centrality (e.g., degree, betweenness) that can determine changes in information flow through local and global brain networks. A threshold of Pearson’s r ≥ 0.85 was used to generate the edges in the networks. In addition, these analyses will be used to determine ‘hub’ regions that may play a disproportionate role in controlling function of the network (Rubinov and Sporns, 2010) and, ultimately, drinking behavior. To identify hub nodes, degree and betweenness scores were ranked separately for male (top 20%) and female (top 10%) water and ethanol drinking mice. Nodes that overlapped between water and ethanol drinking mice for each sex were removed to eliminate nodes that were resistant to ethanol drinking. Overlap in the remaining nodes for degree and betweenness for each sex in the ethanol drinkers were then identified.

### Analysis

Drinking data in high-drinking male and female mice were first analyzed using an unpaired t-test (GraphPad). Next, an Gardner-Altman estimation plot with 5000 bootstrap samples from high-drinking male and female mice were performed using an online analysis tool (www.estimationstats.com). The estimation plots were generated in GraphPad. Pearson’s correlations were used to generate the correlation matrices across the 306 brain regions. The correlation coefficients for individual brain regions in the isocortex, cortical subplate, striatum, and pallidum divisions (*n* = 63 regions) were averaged and analyzed by two-way ANOVA. Two-way ANOVAs were used for analyzing the local transitivity, betweenness, and degree variables. Averaged characteristic path length for each of the 12 major divisions was analyzed by a two-way ANOVA. The threshold for statistical significance was set using an α of 0.05, and Tukey post-hoc test was used for all multiple comparisons.

## Results

### Alcohol Drinking

To generate mice for whole-brain c-Fos mapping, male and female mice were given access to two bottles containing water only (n = 16) or ethanol (20%, v/v; n = 48) with a choice of water in the intermittent access model for 4 weeks. During the last week of 24-h access, there was a range of ethanol drinking across individual C57BL/6J mice with daily ethanol intake averaging between 8.7 and 24.9 g/kg for males and 19.9 and 34.2 g/kg for females (**Figure 1A**). All water drinking mice and mice of both sexes with a heavy ethanol drinking phenotype were included in the whole-brain c-Fos analysis. For these mice, they were given a short (2 h) drinking session in the 5^th^ week before brain extraction 30 min after ethanol availability. The drinking data for these mice on brain extraction day are shown in **Figure 1B**. As expected, there was a range of drinking (2 g/kg to 6 g/kg), and female mice consumed significantly more ethanol on extraction day compared with male mice (**Figure 1C**; *t*_13_ = 2.188, *p* = 0.048). As a complementary analysis, we analyzed the 2 h drinking data using estimation statistics (Ho et al., 2019). The effect size of the unpaired mean difference between ethanol intake in male and female mice is 1.08 (95.0% confidence interval: 0.197, 2.03; **Figure 1D**). The result of a two-sided permutation t-test revealed a significant sex difference in the effect size (*p* = 0.049).

**Figure 1.**
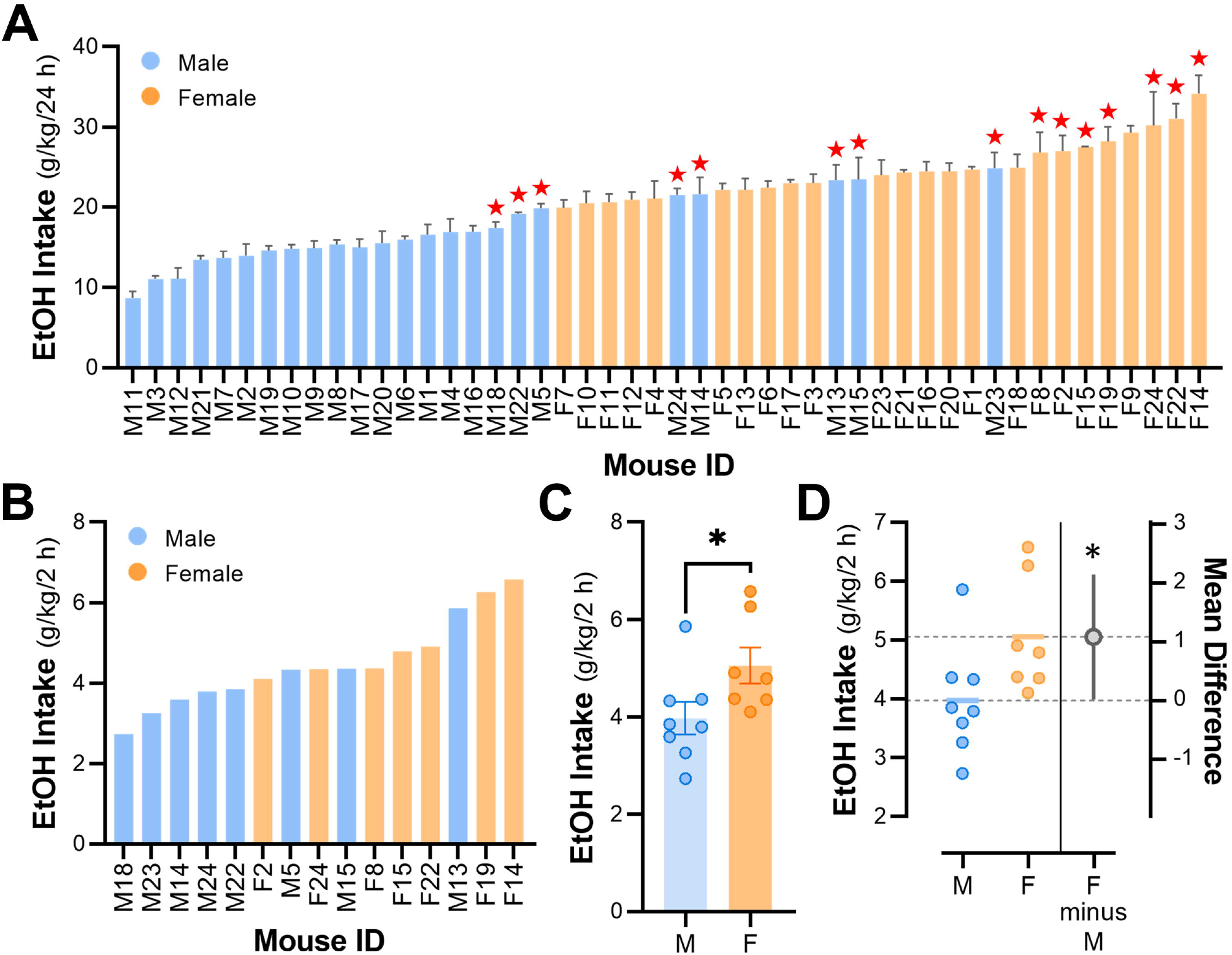
Average daily intake in male and female C57BL/6J mice that were allowed to consume ethanol in the intermittent access, two-bottle choice, model. *(n* = 24 mice/sex). ‘★’ denotes the individual high-drinking male and female mice that were used for whole-brain c-Fos mapping. **(B)** Individual and **(C)** averaged ethanol drinking values on the day of brain extraction. \**p* = 0.048 vs male mice. **(D)** Estimation plot of individual drinking values on extraction day and mean difference for female and male drinking mice (gray dot) with 95% confidence interval shown in the gray bars. \**p* = 0.049 vs 0.

### Correlation Matrices

Representative images of autofluorescence and c-Fos+ cells in a male, ethanol-drinking mouse are shown in **Figure 2**. Raw c-Fos cell count data were generated using BrainQuant3D for each brain region, registered to the Allen Reference Atlas, and then normalized to region volume. Correlation matrices were generated for the 306 regions included in the analyses for each sex and drinking group (**Figure 3A**). What is noticeable from the correlation matrices is global differences in correlation patterns between male and female water-drinking mice, with male mice showing more positive correlations across the entire network. Perhaps more striking is the relative effect of ethanol drinking on regional correlations in female compared to male mice. In females, ethanol drinking increased regions with negative correlations or anti-correlations, whereas ethanol drinking in male mice appeared to strengthen the positive correlations that were more abundant in the water drinking males. As an initial exploration of sex and treatment changes in c-Fos correlations, we calculated the average correlation coefficient within four of the major brain divisions (**Figures 3B**). We observed a significant interaction between sex and treatment group (*F*_1,248_ = 33.88, *p* < 0.001), with post-hoc comparisons revealing higher correlation coefficients in male vs female water drinker (*p* < 0.0001). Moreover, post-hoc analyses revealed that the correlation coefficients were significantly higher in male ethanol drinkers (*p* < 0.01) and reduced in female ethanol drinkers (*p* < 0.0001) compare with their water drinking controls.

**Figure 2.**
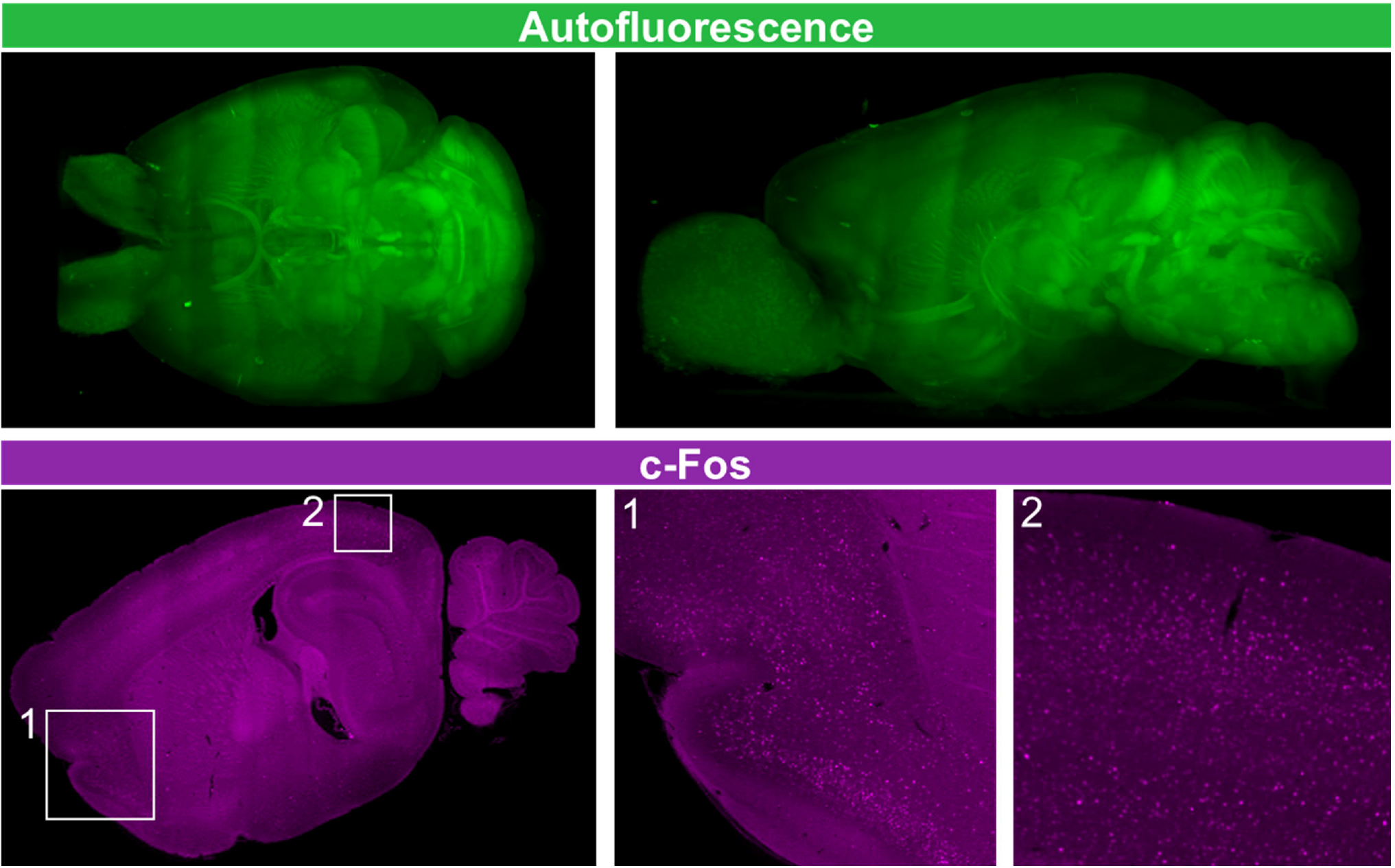
Representative images of autofluorescence and c-Fos+ cells in a male, ethanol-drinking mouse. *Box 1:* c-Fos+ cells in the orbitofrontal and piriform cortices. *Box 2:* c-Fos+ cells in the posterior parietal and visual cortices.

**Figure 3.**
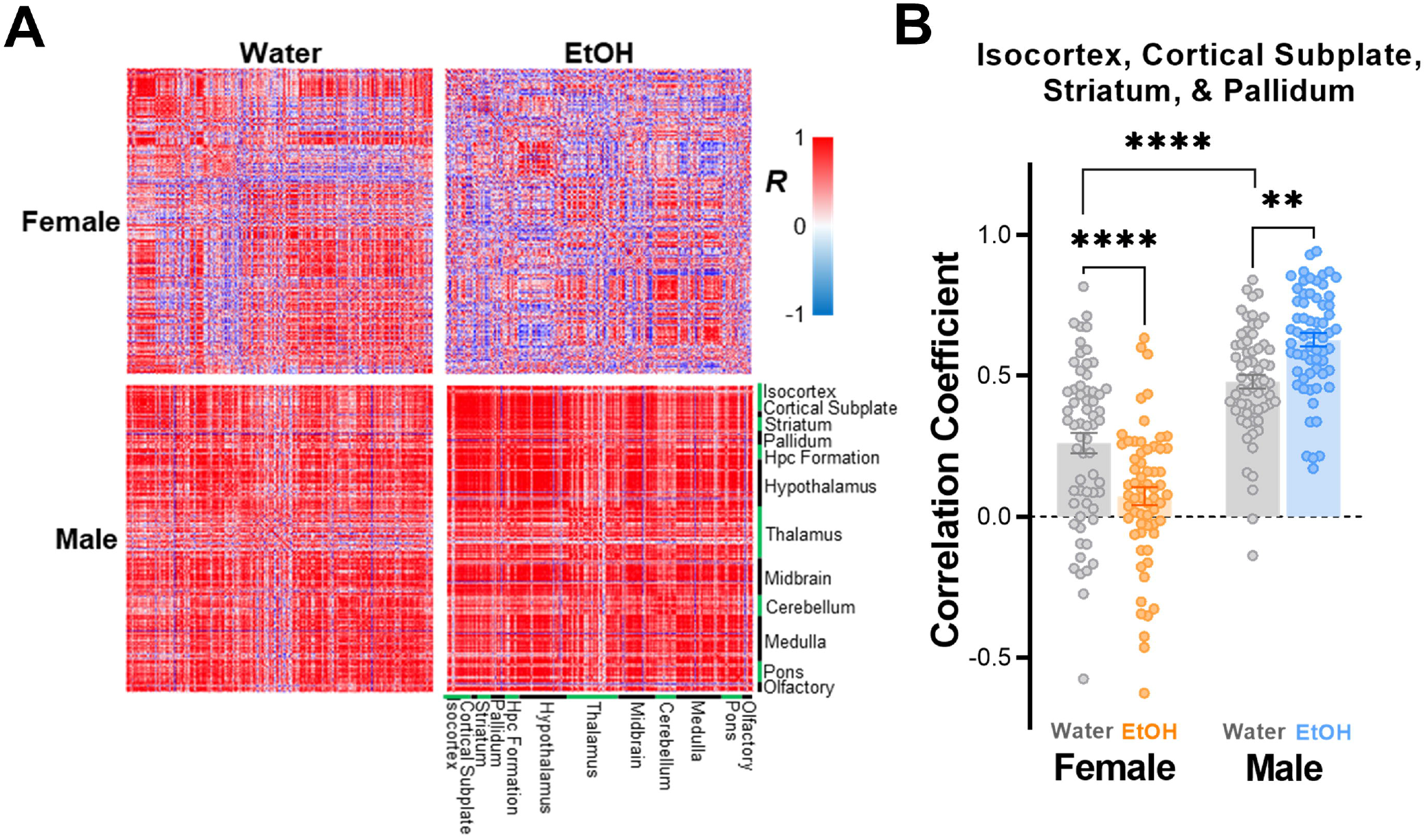
Analysis of whole-brain c-Fos mapping in female and male mice that consumed water or ethanol in the intermittent access model. **(A)** Correlation matrices organized by 12 major divisions of the ARA ontology. **(B)** Average correlation coefficient in 4 major brain divisions (*n* = 63 brain regions). \*\*\*\**p* < 0.0001, female water vs male water and female ethanol; \*\**p* < 0.01, male water vs male ethanol.

### Functional Integration and Segregation Analyses

The networks from the functional connectivity analysis are shown in **Figure 4A**. There was a significant main effect of sex (F_1, 44_ = 9.737, *p* = 0.003), but not drinking solution (F_1, 44_ = 0.074, *p* = 0.786), for characteristic path length, which is a measure of functional integration of distributed brain areas and is the average shortest path length for all pairs of nodes (**Figure 4B**). Similar to the results from the correlation coefficient analysis, there was a significant interaction for transitivity (F_1, 1144_ = 88.24, *p* < 0.0001; **Figure 4C**), which is a measure of functional segregation of the networks. There were also significant interactions for both measures of centrality: degree (F_1, 1220_ = 156.1, *p* < 0.0001; **Figure 4D**) and betweenness (F_1, 1220_ = 18.97, *p* < 0.0001; **Figure 4E**). For all three metrics, post-hoc comparison testing revealed significant differences between water drinking male and female mice (*p* < 0.05). In addition, ethanol drinking significantly affected transitivity and degree in opposite directions for male vs female mice (*p* < 0.01). A similar pattern was observed for betweenness, but the effect of ethanol in males did not reach statistical significance (*p* = 0.057).

**Figure 4.**
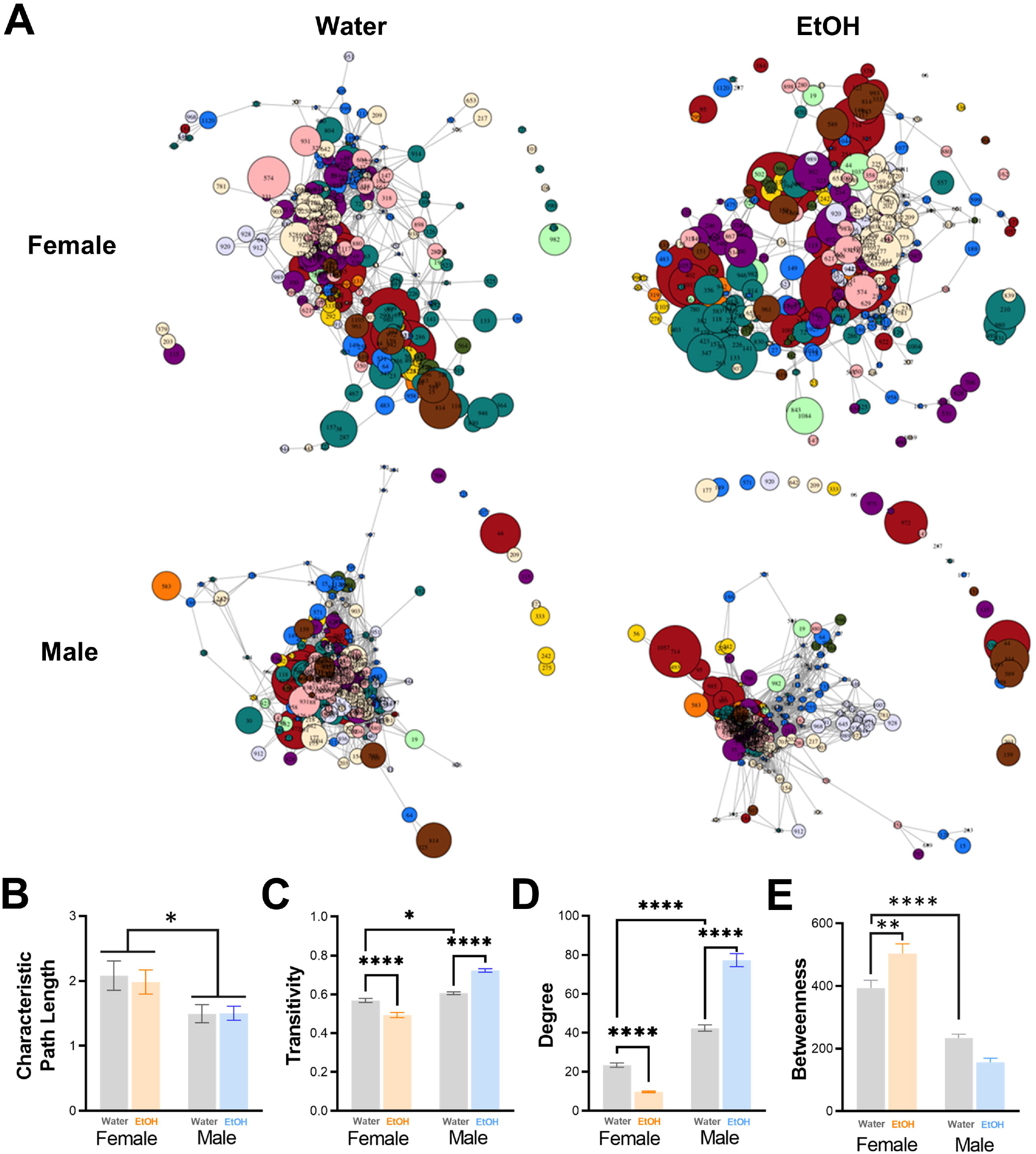
Water and alcohol drinking differentially affect network connectivity in male and female mice. **(A)** Functional connectivity network graphs in male and female water and alcohol drinking mice. Analysis of network metrics for **(B)** characteristic path length (*p < 0.05 vs male mice), **(C)** transitivity (*p < 0.05 vs male water drinking mice, \*\*\*\**p* < 0.0001, water vs ethanol drinking mice), **(D)** degree (\*\*\*\**p* < 0.0001 vs male water drinking mice, \*\*\*\**p* < 0.0001, water vs ethanol drinking mice), and **(E)** betweenness (\*\*\*\**p* < 0.0001 vs male water drinking mice, \*\**p* < 0.01, water vs ethanol drinking female mice).

One goal of these studies is to identify important hub regions that may play a central role in excessive drinking. Using centrality measures, degree and betweenness scores were ranked for male and female water and ethanol drinking mice. Nodes that overlap between water and ethanol drinkers for each sex were removed to eliminate treatment resistant nodes, and duplicate regions for degree and betweenness for each sex in the ethanol drinkers were then identified. This selective ranking of overlapping nodes for each sex resulted in 6 hub nodes in the male ethanol drinkers and 5 hub nodes in the female ethanol drinkers (**Figure 5**).

**Figure 5.**
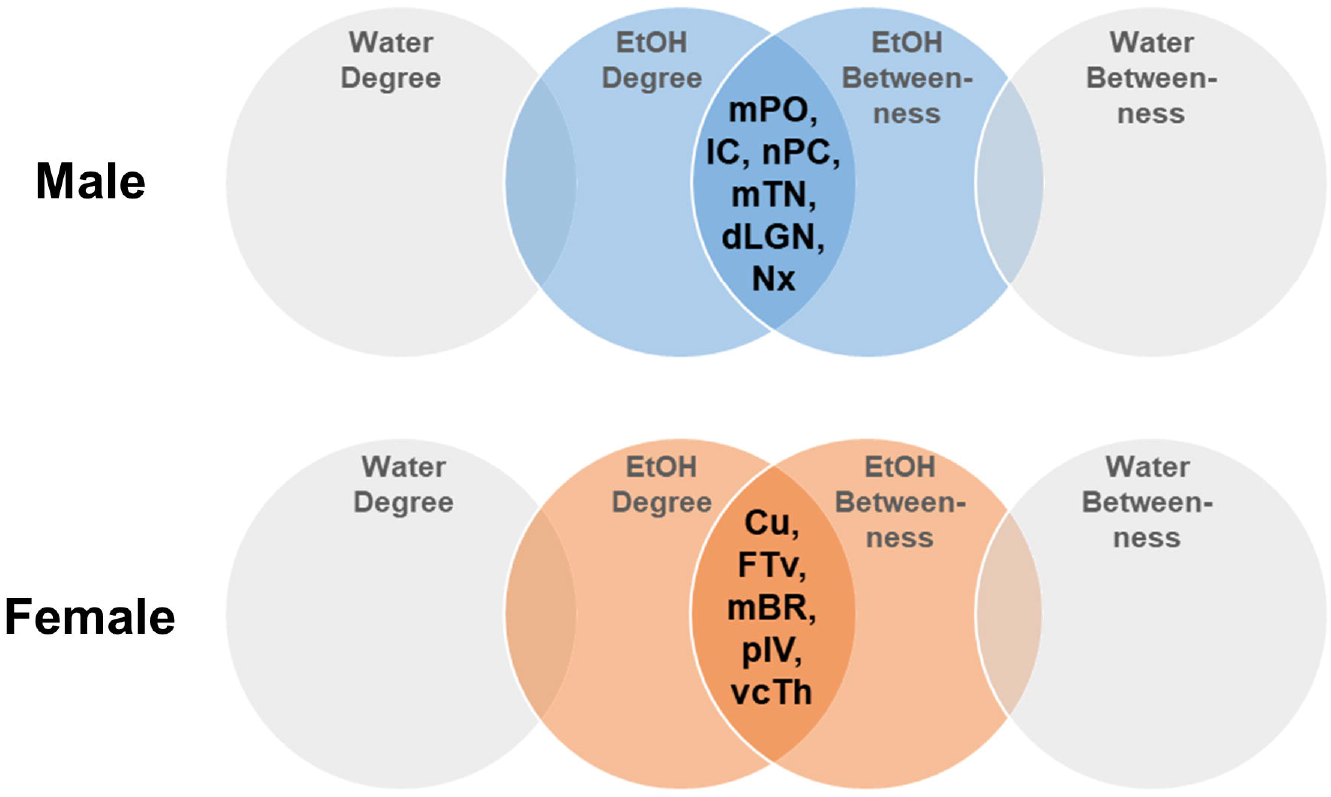
Excessive ethanol drinking engages different hub regions in male and female mice. Venn diagrams showing the overlap in nodes identified using degree and betweenness metrics in male and female water and alcohol drinking mice. *Males:* Medial preoptic nucleus (mPO), Intercalated amygdalar nucleus (IC), Nucleus of the posterior commissure (nPC), Midbrain trigeminal nucleus (mTN), Nucleus x (Nx), and Dorsal part of the lateral geniculate complex (dLGN). *Females:* Culmen (Cu), Folium-tuber vermis (VII) (FTv), Medulla, behavioral state related (mBR), Posterolateral visual area (pIV), Ventral anterior-lateral complex of the thalamus (vcTh).

## Discussion

In the current study, we used whole-brain c-Fos mapping and graph theoretical network approaches to identify mesoscale neural activation induced by excessive ethanol drinking in male and female mice. There were three major findings from our c-Fos mapping study in actively drinking mice. First, our analysis revealed sex differences in the basal global network in water drinkers. Second, there were divergent and sometimes opposing changes in the correlational structure and functional network in excessively drinking male and female mice. Third, using two graph theory metrics for centrality, our analyses identified that excessive drinking engaged different sets of hub regions in male and female mice. Importantly, these nuclei are understudied in the alcohol neuroscience field and will serve as targets in future studies focused on manipulating excessive drinking in the intermittent access drinking model. While clinical and preclinical electrophysiological and imaging studies have examined drinking-induced changes in neural activity after a history of chronic drinking (Cannady et al., 2020; Fang et al., 2021; Nimitvilai et al., 2017; Siciliano et al., 2019; Weber et al., 2014; Zheng et al., 2015), this is the first analysis of the global connectivity changes in male and female mice with a history of excessive drinking that were actively consuming ethanol.

Not surprisingly, we identified sexual dimorphism in the functional connectivity networks across all of the metrics from the graph theory analysis in the water drinking mice. This sexual dimorphism was clearly evident in the correlation matrices and the functional connectivity network graphs. Previous studies have reported anatomical sex differences in human and rodent brains (Goldstein et al., 2001; Qiu et al., 2018; Ruigrok et al., 2014), and analysis of gene expression across the brain also shows sexual dimorphism (Gegenhuber and Tollkuhn, 2020; Liu et al., 2020; Lopes-Ramos et al., 2020; Qiu et al., 2018). Consistent with a structural connectome study across gender in humans (Ingalhalikar et al., 2014), we found that transitivity was higher globally in male water drinking mice compared with female mice. The functional network analysis also revealed sex differences in both measures of centrality and in characteristic path length. Together, these results provide further evidence of sexual dimorphism in global and local connectivity networks in water drinking male and female C57BL/6J mice that reflects some gender-specific outcomes in the human brain connectome.

Perhaps as interesting, excessive ethanol drinking produced robust differences in multiple measures of global and local functional connectivity in male and female mice. Often, the ability of ethanol drinking to change these metrics went in opposite directions for males and females that further separated the sex dimorphism observed in the water drinking mice. In other words, excessive ethanol drinking produced further separation in the functional connectivity networks between male and female mice. Because female mice drink more ethanol than male mice, it is possible that the divergent functional networks are caused by differences in the amount of ethanol consumed. Consistent with previous reports (Bloch et al., 2020; Hwa et al., 2011; Sneddon et al., 2019; Yoneyama et al., 2008), the high-drinking female mice that were included in this c-Fos mapping study consumed more alcohol during the 4^th^ week of drinking compared with the high-drinking male mice. This effect of sex on ethanol intake was also observed during the short drinking episode on the day that brains were extracted and prepared for whole-brain c-Fos mapping. While the amount of ethanol drinking may explain the robust differences between male and female functional networks, we consider this possibility unlikely given the differences in the networks in male and female water drinkers. There was clear separation between the high alcohol drinking males and females when given 24 h of ethanol access, there was less clear separation across sexes on the last drinking session. On the day of brain extraction, female mice consumed ~1.1 g/kg more than male mice in the 2 h drinking session. However, male mice still consumed a relatively large amount of ethanol in the short drinking session (i.e., ~4 g/kg). Although we did not measure blood ethanol concentrations (BECs) at the time of sacrifice, we would expect that BECs would be above a binge level of intoxication (>80 mg/dl) in both sexes. We hypothesize that the divergent changes in the functional connectivity networks produced by excessive ethanol drinking are related to sexual dimorphism in neuroanatomy and gene expression rather than the amount of lifetime ethanol consumption. Additional whole-brain c-Fos mapping analysis in the low-ethanol drinking male and female mice will help to provide some insight into this unanswered question.

In addition to the divergent and opposing networks produced by excessive ethanol drinking, it is not surprising that the two measures of centrality also identified different hub regions in the male and female ethanol drinking mice. In the male drinking mice, hub nuclei were identified in the thalamus, hypothalamus, midbrain, and medulla. In contrast, a unique set of hub nuclei were identified in the cerebellum, medulla, isocortex, and thalamus of excessively drinking female mice. A strength of this unbiased, whole-brain network analysis is to reveal novel and understudied brain regions that may be important for driving excessive ethanol drinking. Virtually nothing is known about the actions on ethanol on the identified hub nuclei (e.g., midbrain trigeminal nucleus) or if they control voluntary or excessive ethanol drinking. One study reported that the midbrain trigeminal nucleus regulates food and water in mice intake possibly through influencing hypothalamic orexinergic signaling (Yokoyama et al., 2013). Recent evidence suggests that midbrain structures can control compulsive drinking (Siciliano et al., 2019), and the orexin system has been linked to the motivation effects of alcohol (Anderson et al., 2018). Thus, the midbrain trigeminal nucleus may also regulate excessive alcohol drinking in male, but not female mice. Although studies are required to determine the role of these understudied regions, such as the midbrain trigeminal nucleus, in regulation of ethanol intake, the analysis revealed a set of potentially critical and sexually unique nuclei that can be target in future studies in male and female mice.

In conclusion, the findings from these studies revealed sexually dimorphic responses in whole-brain c-Fos activity mapping in water drinking male and female mice and robust and divergent shifts in the functional connectivity networks in mice that consumed excessive amounts of ethanol over a 4-week period. In addition to the effects of sex and ethanol drinking on functional segregation and integration, these analyses also identified unique and understudied regions that may play a critical role in controlling excessive ethanol drinking in male and female mice. Combined with the findings from another study measuring network changes in functional connectivity in ethanol dependent mice (Kimbrough et al., 2020; Simpson et al., 2021), unbiased whole-brain immediate early gene mapping, light sheet imaging, and advanced functional connectivity network analysis is a promising tool to generate new hypotheses and identify potentially important regions that control AUD-related behaviors, such as harmful ethanol intake.

## Notes

**Conflict of Interest:** Authors report no conflict of interest.

**Funding Sources**: These studies were supported by NIH grants AA020930 (PJM), AA023288 (PJM), and a pilot project funded by the Charleston Alcohol Research Center (AA010761). In addition, these studies were supported by the Department of Veterans Affairs, Veterans Health Administration, Office of Research Development (BLR&D) and VA grant BX000798 (PJS).

### Competing Interest Statement

The authors have declared no competing interest.

